# Molecular physiology of chemical defenses in a poison frog

**DOI:** 10.1101/591115

**Authors:** Stephanie N. Caty, Aurora Alvarez-Buylla, Gary D. Byrd, Charles Vidoudez, Alexandre B. Roland, Elicio E. Tapia, Bogdan Budnik, Sunia A. Trauger, Luis A. Coloma, Lauren A. O’Connell

**Author notes:** To whom correspondence should be addressed: Lauren A. O’Connell, Department of Biology, Stanford University, 371 Serra Mall, Stanford, CA 94305.

## Abstract

Poison frogs sequester small molecule lipophilic alkaloids from their diet of leaf litter arthropods for use as chemical defenses against predation. Although the dietary acquisition of chemical defenses in poison frogs is well-documented, the physiological mechanisms of alkaloid sequestration has not been investigated. Here, we used RNA sequencing and proteomics to determine how alkaloids impact mRNA or protein abundance in the Little Devil Frog (*Oophaga sylvatica*) and compared wild caught chemically defended frogs to laboratory frogs raised on an alkaloid-free diet. To understand how poison frogs move alkaloids from their diet to their skin granular glands, we focused on measuring gene expression in the intestines, skin, and liver. Across these tissues, we found many differentially expressed transcripts involved in small molecule transport and metabolism, as well as sodium channels and other ion pumps. We then used proteomic approaches to quantify plasma proteins, where we found several protein abundance differences between wild and laboratory frogs, including the amphibian neurotoxin binding protein saxiphilin. Finally, because many blood proteins are synthesized in the liver, we used thermal proteome profiling as an untargeted screen for soluble proteins that bind the alkaloid decahydroquinoline. Using this approach, we identified several candidate proteins that interact with this alkaloid, including saxiphilin. These transcript and protein abundance patterns suggest the presence of alkaloids influences frog physiology and that small molecule transport proteins may be involved in toxin bioaccumulation in dendrobatid poison frogs.

**Resumen:** Las ranas venenosas obtienen moléculas lipofílicas a partir de su dieta de artrópodos que luego usan como una defensa química contra depredadores. Mientras que la acumulación de toxinas dietéticas ha sido bien documentada, el mecanismo fisiológico de obtención de alcaloides no ha sido investigado. En este estudio usamos secuenciación de RNA y proteómica para determinar cómo la presencia de alcaloides afecta la abundancia de mRNA y proteínas en ranas diablito (*Oophaga sylvatica*) silvestres con defensas químicas en comparación a ranas diablito criadas en laboratorio con una dieta sin alcaloides. Para entender cómo las ranas venenosas mueven los alcaloides de su dieta a las glándulas granulares en su piel, nos enfocamos en medir la expresión de genes en tres tejidos: intestinos, piel e hígado. En estos tejidos, encontramos varios transcriptomas regulados diferencialmente que tienen actividades involucradas con el transporte y metabolismo de pequeñas moléculas, además de canales de sodio y bombas de iones. Luego usamos métodos proteómicos para cuantificar proteínas en plasma, donde encontramos varias diferencias en abundancia de proteínas entre las ranas silvestres y de laboratorio, incluyendo la proteína anfibia de fijación de toxinas, saxifilina. Finalmente, debido a que muchas proteínas encontradas en la sangre se sintetizan en el hígado, usamos la técnica de perfilación proteómica termal para seleccionar imparcialmente las proteínas solubles que fijan el alcaloide decahydroquinolina. Usando este método, identificamos varias posibles proteínas que interactúan con este alcaloide, incluyendo saxifilina. Estos patrones de cambios en abundancia de transcriptomas y proteínas en ranas con y sin defensas químicas sugieren que la presencia de alcaloides influye en la fisiología de las ranas y que moléculas proteicas pequeñas de transporte podrían estar involucradas en la bioacumulación de toxinas en ranas venenosas dendrobátidos.

**Summary Statement:** Chemically defended wild poison frogs have gene expression and protein abundance differences across several tissue systems compared to poison frogs reared on an alkaloid-free diet.

## Introduction

Many animals, from microbes to vertebrates, have chemical defenses to guard against predation. Although microbes and plants produce secondary metabolites that can serve as toxins (Burja et al., 2001; Gershenzon, 1999), most animals cannot synthesize small molecule toxins themselves. While some animals like snakes and arachnids produce genetically encoded peptide toxins, other vertebrate animals, including some amphibians, reptiles, and birds, carry small molecule toxins that are acquired from dietary sources (Savitzky et al., 2012a). Invertebrates can also sequester small molecule toxins from their diet and this process has been documented in beetles, moths and butterflies (Trigo, 2010; Petschenka and Agrawal, 2016) Many decades of research has focused on cataloging the small molecules sequestered by vertebrates and documenting the trophic relationships of these chemically defended animals and their prey (Savitzky et al., 2012b).

The ability to sequester small molecule toxins from dietary sources has evolved many times in frogs, including Neotropical dendrobatids and bufonids, Malagasy mantellids, Australian myobatrachids (*Pseudophryne*) and Cuban eleutherodactylids (Saporito et al., 2011). These frogs uptake lipophilic alkaloids from arthropod prey items, store them in skin granular glands, and secrete them as a defensive response (Santos et al., 2016). In order to utilize diet-derived toxins, frogs may be resistant to alkaloids they are accumulating through mutations in target proteins. Many alkaloids target voltage-gated sodium channels and nicotinic acetylcholine receptors in the nervous system and some studies have found ion channel mutations that confer toxin resistance in poison frogs (Santos et al., 2016; Tarvin et al., 2016; Tarvin et al., 2017; Wang and Wang, 1998). Detailed information on binding properties of a wide range of poison frog alkaloids on many different ion channel types is lacking as early studies focused on a few types of alkaloids and ion channels. Despite the progress on understanding the evolution of autoresistance and chemical defenses in poison frogs, no study has examined other molecular aspects of physiology that allows toxin sequestration.

To accomplish the sequestration of alkaloids, poison frogs selectively move these small molecules from the gastrointestinal tract to the skin glands for storage. In many animals, toxic compounds are unable to pass through the intestinal wall for absorption due to a series of membrane transporters in the gut epithelium. These transporters bind compounds and move them either into circulation or back into the intestinal lumen for excretion (Steffansen et al., 2004). Once absorbed in the intestines, lipophilic compounds can be circulated by a number of different mechanisms, including binding by carrier proteins in the blood circulation or within lipoprotein particles like chylomicrons in the lymph system (Trevaskis et al., 2008). In many animals, dietary compounds in the blood circulation are metabolized by the liver and there is some evidence that poison frogs can metabolize dietary alkaloids. Daly and colleagues suggested that some (but not all) dendrobatid poison frog species can metabolize pumiliotoxin **251D** into allopumiliotoxin **267A** (Daly et al., 2003), although the mechanisms underlying this observation remain unknown. Clearly coordination across several tissue systems is required for alkaloid sequestration in poison frogs, but the physiological basis for alkaloid bioaccumulation remains unexplored.

The goal of this present study was to better understand the physiology of acquired chemical defenses in dendrobatid poison frogs. In surveying the alkaloid profiles of wild caught Little Devil frogs (*Oophaga sylvatica,* Family Dendrobatidae) and juveniles captured in the wild but raised in laboratory on an alkaloid-free diet, we had an opportunity to examine physiological differences associated with chemical defenses. We compared gene expression in the intestines, liver, and skin, as well as plasma protein abundance in wild caught chemically defended frogs and laboratory frogs raised on an alkaloid-free diet. To further functionally identify proteins that may bind toxins for transport, we performed a thermal proteome profiling assay using liver protein lysate from a laboratory-reared frog. As toxins are acquired from the diet and transported to the skin granular glands for storage, we expected that the presence of toxins would cause an increase in expression of genes associated with small molecule transport or metabolism.

## Materials and Methods

### Field collection of *Oophaga sylvatica*

Little Devil (or Diablito) frogs [*Oophaga sylvatica*, (Funkhouser) N=13] were collected during the day near the Otokiki Reserve (GPS coordinate: latitude 0.91075, longitude −78.57555, altitude 704m) maintained by WIKIRI/Fundación Otonga in northwestern Ecuador in July 2013 (Roland et al., 2017). Frogs were individually stored in plastic bags with air and vegetation for 2–5 hours. In the evening the same day of capture, adult frogs (over one year of age; four females and four males) were anesthetized with a topical application of 20% benzocaine to the ventral belly and euthanized by cervical transection. Several juvenile frogs (sex unknown, N=5) were transported to Centro Jambatu de Investigación y Conservación de Anfibios in Quito, Ecuador (CJ), where they were individually housed and fed a diet of crickets and fruit flies for 6 months before being sacrificed as adults as described above. For each individual, the intestines and liver were dissected and stored in RNALater (Life Technologies, Carlsbad, CA, USA). Half of the dorsal skin was stored in RNALater while the other half was stored in methanol for later chemical analysis. Remaining frog tissues were preserved in 100% ethanol and deposited in the amphibian collection of CJ. Additional frogs (N=6) were collected in January 2014 in the Otokiki Reserve and, along with some captive adults raised at CJ (N=6), were treated as described above with one additional procedure: after euthanasia, trunk blood was collected into a heparinized microcentrifuge tube. Blood was centrifuged and plasma was isolated into a fresh tube and flash frozen in liquid nitrogen. Collections and exportation of specimens were done under permits (001-13 IC-FAU-DNB/MA, CITES 32/VS and 37/VS) issued by the Ministerio de Ambiente de Ecuador. Sample sizes were chosen based on typical samples sizes in the literature as group variance was unknown prior to the experiments. Sample sizes were minimized to reduce impact on wild *O. sylvatica* populations. The Institutional Animal Care and Use Committee of Harvard University approved all procedures (Protocol 15-02-233).

### Gas chromatography / mass spectrometry (GC/MS)

Frog dorsal skin samples (N=5 wild caught and N=3 laboratory raised frogs) were used to characterize alkaloid profiles as previously described (McGugan et al., 2016). As an internal standard, 25 μg of D3-nicotine in methanol (Sigma-Aldrich, St. Louis, MO) was added to each sample prior to alkaloid extraction. After extraction in methanol, the samples were evaporated to dryness under nitrogen gas and reconstituted in 0.5 mL of methanol by vortexing. The samples were transferred to a microcentrifuge tube and spun at 12000 rpm for 10 min. A 200 μL aliquot of the supernatant was transferred to a 0.3 mL glass insert in an amber sample vial and stored at −20°C until analysis.

GC/MS parameters were conducted as previously described (McGugan et al., 2016), which is based on a slight modification of the method reported by Saporito and colleagues (Saporito et al., 2010b). Briefly, analyses were performed on a Waters Quattro Micro system (Beverly, MA) with an Agilent 6890N GC (Palo Alto, CA). Each total ion chromatogram was reviewed and alkaloid responses were characterized from their mass spectra and retention times. The extensive database developed by Daly and colleagues (Daly et al., 2005) was used to identify alkaloids by matching major mass spectrum peaks and relative retention times using D3-nicotine as a reference. AMDIS (NIST) was used in conjunction with the database to identify candidate alkaloids peak. Those peaks were then manually inspected and the confirmed alkaloids peaks were then integrated. Unidentifiable compounds were removed from data analysis. The relative abundance values were normalized by nicotine and were visualized using the heatmap.2 R function in the gplots package. A representative extracted ion chromatograms for each group was normalized to the same scale to create a visual overview. All GC/MS data is available at DataDryad (submission pending).

### Reference transcriptome

To build a reference transcriptome for gene expression analyses, we sequenced tissues from a single *O. sylvatica*. We isolated intestines, liver and dorsal skin from a single laboratory-bred *O. sylvatica* frog (Indoor Ecosystems Whitehouse, Ohio, USA) for the purpose of *de novo* transcriptome assembly. Flash frozen tissue samples were placed in Trizol (Life Technologies, Grand Island, NY) where RNA was extracted according to manufacturer instructions. Polyadenylated RNA was isolated from each sample using the NEXTflex PolyA Bead kit (Bioo Scientific, Austin, TX, USA) according to manufacturer instructions. Lack of contaminating ribosomal RNA was confirmed using the Agilent 2100 Bioanalyzer. Strand specific libraries for each sample (intestines, liver, and skin) were prepared using the dUTP NEXTflex RNAseq kit (Bioo Scientific), which includes a magnetic bead-based size selection. Libraries were pooled in equimolar amounts after library quantification using both quantitative PCR with the KAPA Library Quantification Kit (KAPA Biosystems, Wilmington, MA, USA) and the fluorometric Qubit dsDNA high sensitivity assay kit (Life Technologies), both according to manufacturer instructions.

Libraries were sequenced on an Illumina HiSeq 2500 over a full flow cell to obtain 433,951,467 paired-end 250 bp reads. We first corrected errors in the Illumina reads using Rcorrector ((Song and Florea, 2015); parameters: run_rcorrector.pl -k 31) and then applied quality and adaptor trimming using Trim Galore! (http://www.bioinformatics.babraham.ac.uk/projects/trim_galore/; parameters: trim_galore -- paired --phred33 --length 36 −q 5 --stringency 5 --illumina −e 0.1). We used Trinity to *de novo* assemble the *O. sylvatica* transcriptome ((Grabherr et al., 2011), parameters: --seqType fq -- SS_lib_type RF --normalize_reads). The Trinity assembly produced 2,111,230 contigs (N50: 738 bp). To reduce redundancy in the assembly, we ran cd-hit-est ((Fu et al., 2012); parameters: -c 0.98) resulting in 1,596,373 contigs (N50: 789 bp). For the purpose of our current study, we were specifically interested in putative protein coding transcripts to gain insight into the physiological mechanisms of toxin sequestration. We used blastx to compare contigs in the assembly to proteins in the Uniprot Swiss-Prot database (e-value threshold of 1e-5) and retained 241,116 contigs (N50: 2169 bp) with homology to annotated proteins. As we were particularly interested in frog transcripts, we removed contigs that had homology to non-vertebrate proteins in the Uniprot Swiss-Prot database, including microbes, nematodes, and arthropods. These contaminants that likely represent parasites, pathogens, and prey items represented roughly 20% of the contigs in the draft assembly. Our final *O. sylvatica* draft assembly contained 194,309 contigs (N50=2280 bp). We then assessed the completeness of this filtered assembly by examining vertebrate ortholog representation using BUSCO (Benchmarking Universal Single-Copy Orthologs) (Simão et al., 2015). BUSCO estimated the completeness of our assembly at 84% (700) complete single-copy BUSCOs, with 61% (1862) duplicated BUSCOs, 5.9% (181) fragmented BUSCOS and 9.2% (280) missing BUSCOs out of 3023 total BUSCO groups searched.

We annotated the transcriptome using Trinotate (Bryant et al., 2017) and used this final assembly for downstream gene expression analyses. We used TransDecoder (https://github.com/TransDecoder/TransDecoder/wiki) to generate a peptide file populated by the longest open reading frame for each contig, which was then annotated using blastp to the UniProt database and hmmscan to the PFam-A database. The results of these database queries were loaded into an SQlite database.

### RNA sequencing and analysis

The alkaloid differences between wild and laboratory raised frogs presented a unique opportunity to examine gene expression differences associated with chemical defense (see results). We focused on the liver and skin, as alkaloid compounds have been detected in these tissues in other poison frog species previously (Saporito et al., 2011). We also analyzed gene expression in the intestines as a likely location of alkaloid absorption (Estudante et al., 2013). We isolated RNA from the intestines, liver, and dorsal skin of the laboratory frogs raised on an alkaloid-free diet and wild caught frogs, including the same individuals used for the GC/MS analysis. Illumina libraries were constructed as described above and sequenced on an Illumina HiSeq 2000 over 6 lanes to obtain 910,220,544 paired-end 100bp reads. All Illumina data is available on the Sequence Read Archive (submission pending).

We first applied quality and adaptor trimming using Trim Galore! (http://www.bioinformatics.babraham.ac.uk/projects/trim_galore/; parameters: trim_galore -- paired --phred33 --length 36 −q 5 --stringency 5 --illumina −e 0.1). We analyzed tissues separately because some individuals were removed from specific tissue comparisons because these samples had fewer than 6 million reads. We aligned the reads to the *O. sylvatica* reference transcriptome (see above) using kallisto (Bray et al., 2016). Differences in gene expression levels between samples were analyzed using edgeR (Robinson et al., 2010) and DESeq2 (Love et al., 2014) within the *in silico* Trinity pipeline (p < 0.05 FDR, 4-fold change). Gene expression differences were considered significant at p<0.05 with false discovery correction applied (default for DESeq2 is Benjamin-Hochberg correction). We planned on retaining significant differences in transcript abundance if this difference was present in both the edgeR and DESeq2 analyses. However, all transcripts with significant differences in DESeq2 were present in the edgeR significant gene set, but not vice versa, where there were 2-3 times the number of significant genes in the edgeR analysis. Thus, only DESeq2 results are presented in the manuscript to be more conservative. All gene expression results from the DESeq2 analysis were loaded into the SQlite database created from the Trinotate pipeline. The R package GOseq (Young et al., 2010) (version 1.34.1) was used to examine gene ontology enrichment in transcript expression for each tissue within the *in silico* Trinity pipeline. A subset of differentially expressed genes were visualized using boxplots of the TMM normalized expression data output from Trinity, which were made with the R package *ggplot2* (R version 3.4.3).

### Plasma proteomics

As alkaloids can potentially be transported through the blood circulation to the skin glands for storage, we quantified plasma protein abundance in chemically defended wild caught frogs and laboratory frogs reared on an alkaloid-free diet. To have enough protein concentration for tandem mass tag labeling and quantitative proteomics, plasma was pooled for two frogs to create three biological replicates per group. Tandem Mass Tag (TMT; Thermo Scientific, Waltham, MA, USA) labeling was performed according to manufacturer’s instructions. TMT is a chemical label that allows the quantification of proteins from pooled samples by adding slight variations to the molecular mass of proteins. Samples were off-line fractionated prior to liquid chromatography tandem mass spectrometry (LC-MS/MS) analysis performed on a LTQ Orbitrap Elite (Thermo Scientific) equipped with a Waters NanoAcquity high performance liquid chromatography (HPLC) pump (Milford, MA, USA). Peptides were separated onto a 100 μm inner diameter microcapillary trapping column packed first with approximately 5 cm of C18 Reprosil resin (5 μm, 100 Å, Dr. Maisch GmbH, Germany) followed by analytical column ~20 cm of Reprosil resin (1.8 μm, 200 Å, Dr. Maisch GmbH, Germany). Separation was achieved through applying a gradient from 5–27% ACN in 0.1% formic acid over 90 min at 200 nl min−1. Electrospray ionization was enabled through applying a voltage of 1.8 kV using a home-made electrode junction at the end of the microcapillary column and sprayed from fused silica pico tips (New Objective, Woburn, MA, USA). The LTQ Orbitrap Elite was operated in data-dependent mode for the mass spectrometry methods. The mass spectrometry survey scan was performed in the Orbitrap in the range of 395–1,800 m/z at a resolution of 6 × 104, followed by the selection of the twenty most intense ions (TOP20) for CID-MS2 fragmentation in the ion trap using a precursor isolation width window of 2 m/z, AGC setting of 10,000, and a maximum ion accumulation of 200 ms. Singly charged ion species were not subjected to CID fragmentation. Normalized collision energy was set to 35 V and an activation time of 10 ms. Ions in a 10 ppm m/z window around ions selected for MS2 were excluded from further selection for fragmentation for 60 s. The same TOP20 ions were subjected to HCD MS2 event in Orbitrap part of the instrument. The fragment ion isolation width was set to 0.7 m/z, AGC was set to 50,000, the maximum ion time was 200 ms, normalized collision energy was set to 27V and an activation time of 1 ms for each HCD MS2 scan. All LC/MS-MS data is available at DataDryad (submission pending).

Raw data were analyzed using Proteome Discoverer 2.1.0.81 software (Thermo Scientific). Assignment of MS/MS spectra was performed using the Sequest HT algorithm by searching the data against the *O. sylvatica* protein sequence database and other known contaminants such as human keratins and common laboratory contaminants. To create an mRNA-based protein reference database for peptide matching, we used the PHROG tool (**P**roteomic Reference with **H**eterogeneous **R**NA **O**mitting the **G**enome; (Wühr et al., 2014); http://kirschner.med.harvard.edu/tools/mz_ref_db.html). We used as input the Trinity fasta file that had cd-hit-est applied (see above) containing hits to the Uniprot Swiss-Prot database (both vertebrates, invertebrates, and other potential contaminants like microbes). Briefly, this tool identifies ORF frames, frameshift corrections and annotation using blastx to a deuterostome database. The longest peptide is retained and database redundancy is removed with cd-hit. This output is then used as the protein reference for peptide matching. The reference proteome is available at DataDryad (submission pending).

Sequest HT searches were performed using a 20 ppm precursor ion tolerance and requiring each peptides N-/C termini to adhere with Trypsin protease specificity, while allowing up to two missed cleavages. 6-plex TMT tags on peptide N termini and lysine residues (+229.162932 Da) was set as static modifications while methionine oxidation (+15.99492 Da) was set as variable modification. A MS2 spectra assignment false discovery rate (FDR) of 1% on protein level was achieved by applying the target-decoy database search. Filtering was performed using Percolator (Kall et al., 2008). For quantification, a 0.02 m/z window centered on the theoretical m/z value of each the six reporter ions and the intensity of the signal closest to the theoretical m/z value was recorded. Reporter ion intensities were exported in result file of Proteome Discoverer 2.1 search engine as an excel table. The total signal intensity across all peptides quantified was summed for each TMT channel, and all intensity values were adjusted to account for potentially uneven TMT labeling and/or sample handling variance for each labeled channel. Sequest found 453 protein matches within the *O. sylvatica* protein database across all six plasma samples. We omitted from our analysis proteins that had only one peptide match. The data was then imported into Perseus proteomics analysis software (Tyanova et al., 2016) and a t-test was used to examine differences between groups.

### Thermal proteome profiling

A single laboratory-reared *O. sylvatica* frog was purchased from the Indoor Ecosystems (Whitehouse, Ohio, USA) and sacrificed as described above to conduct the thermal proteome profiling experiment (Franken et al., 2015). This assay relies on thermodynamic principles of protein-ligand binding, where a protein that has bound its small molecule target has increased thermal stability, a shift detectable by tandem mass spectrometry. The liver was dissected, flash frozen in liquid nitrogen, and stored at −80°C for 2 weeks. On the day of processing, the liver was crushed in liquid nitrogen with a mortar and pestle. Powdered tissue was then scraped into a Dounce tissue homogenizer and 1.2 mL of ice-cold 1X PBS supplemented with EDTA-free cOmplete protease inhibitor cocktail (Roche, Indianapolis, IN, USA) was added. After the tissue powder was homogenized with 10 plunges of the homogenizer, the cell suspension was transferred to a chilled microcentrifuge tube. For cell lysis, the sample was placed in liquid nitrogen for 1 min followed by a brief thaw at room temperature, and then this cycle was repeated twice. Cell lysate was separated from cellular debris by spinning 20,000xg for 20 min at 4°C. The sample was then divided in half (~400 uL) and either 4 uL of 10mM DHQ (decahydroquinoline, Sigma) in DMSO (100 mM final concentration) or 4 uL of DMSO (vehicle treatment) was added. Lysate was incubated for 30 min at room temperature and then 35 uL of each samples was aliquoted into each of 10 PCR tubes. PCR tubes were heated at different temperatures (67°C, 64.3°C, 58.5°C, 56.2°C, 53°C, 50°C, 47.4°C, 45.1°C, 39.9°C, 37°C) for 3 min followed immediately by 2 min room temperature incubation. Immediately after this incubation, samples were flash frozen in liquid nitrogen and stored at −80°C overnight. The following morning, samples were thawed on ice and transferred to 0.2 mL polycarbonate thick-wall tubes (Beckman Coulter, Danvers, MA, USA) and spun at 100,000xg for 20m at 4°C in Optima Max ultracentrifuge (Beckman Coulter). Tubes were carefully removed from the ultracentrifuge with forceps and 30 uL of supernatant was removed to a PCR tube with every care taken to avoid the protein pellet. Soluble protein lysate was flash frozen in liquid nitrogen and stored at −80°C until downstream processing. Samples were then TMT labeled and analyzed using LC MS/MS as described above and following (Franken et al., 2015). Data was analyzed using the TPP package version 3.10.1 in R version 3.5.2, which fits protein abundances to a curve and generates p-values for proteins with a significant thermal shift when incubated with the ligand of interest compared to the vehicle control. Thermal shift graphs were output using the TPP package and customized using ggplot in R. All LC/MS-MS data is available on DataDryad (submission pending).

## Results

### Alkaloid profiles of wild and captive frogs

Wild caught frogs contain many alkaloid chemicals (Fig 1), the most abundant of which include four decahydroquinolines (**219A**, **219C**, **223Q**, **251A**), several 3,5,-disubstituted indolizidines (**223AB** and **249A**), a 5,8-disubstituted indolizidine (**235B**), two lehmizidines (**277A** and **275A**), and an unclassified alkaloid with unknown structure (**283C**). Control frogs captured as juveniles and housed in laboratory for six months on a diet of fruit flies and crickets contained only trace or undetectable quantities of alkaloids compared to wild caught frogs. The major compounds detected in laboratory-reared frogs were nicotine-D3 (spiked internal reference), benzocaine (anesthetic), and cholesterol remaining from the extraction process.

**Figure 1.**
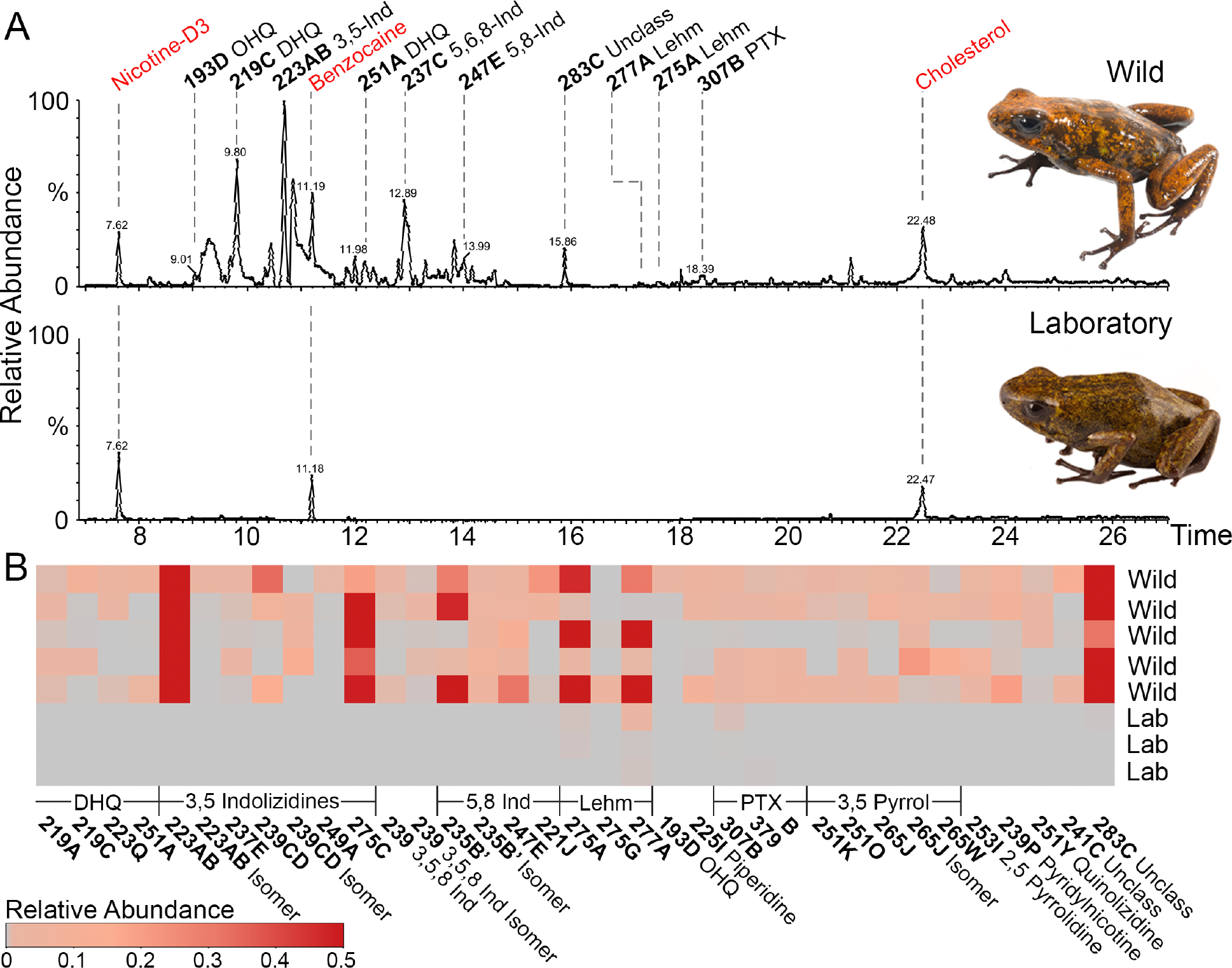
Skin alkaloid abundance differs between wild caught and laboratory-reared frogs from the Otokiki population of *Oophaga sylvatica*. Alkaloid profiles assayed by gas chromatography / mass spectrometry (GC/MS) show chemical defenses in wild caught (top, N=5) and laboratory (bottom, N=3) frogs; this experiment was performed once. **(a)** Ion chromatograms show small molecule peaks (labels at the top); compounds common to both groups are highlighted in red. **(b)** Heatmap shows relative abundance of alkaloid compounds in each frog, highlighting the zero or trace alkaloid quantities in laboratory lab-reared frogs compared to wild caught frogs. Abbreviations: DHQ, decahydroquinoline; 5,8 Ind, 5,8, disubstituted indolizidines; Lehm, lehmizidine; PTX, pumiliotoxin; 3,5 Pyrrol, 3,5 pyrrolizidine.

### Gene expression changes associated with chemical defenses

We next examined how chemical defenses acquired through differences in diet impact physiology by comparing gene expression in several tissues of chemically defended wild caught frogs and laboratory frogs reared on an alkaloid-free diet. As alkaloids are acquired from the diet and stored in the skin granular glands, we used RNA sequencing to measure gene expression in the intestines, skin, and liver.

#### Intestines

Comparing the intestinal gene expression profiles of wild caught and laboratory-raised frogs, 47 transcripts were differentially expressed between the two groups (Fig 2a). Chemically defended wild frogs had increased expression of 9 transcripts and decreased expression of 38 transcripts compared to the laboratory-reared frogs (See Supplementary Excel File). Gene ontology (GO) analyses showed enrichment of several interesting processes: molecular function included ligand gated sodium channel activity (p = 0.002), sodium:chloride symporter activity (p = 0.011), and sodium channel activity (p = 0.029). Enriched cellular compartments included sodium channel complex (p = 0.003), extracellular region part (p = 0.004), and apical plasma membrane (p = 0.004). There were many biological processes enriched in the differentially expressed genes, with the top two being sodium ion transmembrane transport (p = 0.0007) and small molecule metabolic process (p = 0.002). Given the prominence of sodium channels and small molecule metabolism in the GO processes, we further explored the expression of transcripts whose proteins are involved in small molecule transport (sodium channels and solute carrier proteins) and metabolism (cytochrome p450s) (Fig 2b), which mostly show a decrease in expression in wild caught animals compared to animals reared on an alkaloid-free diet.

**Figure 2.**
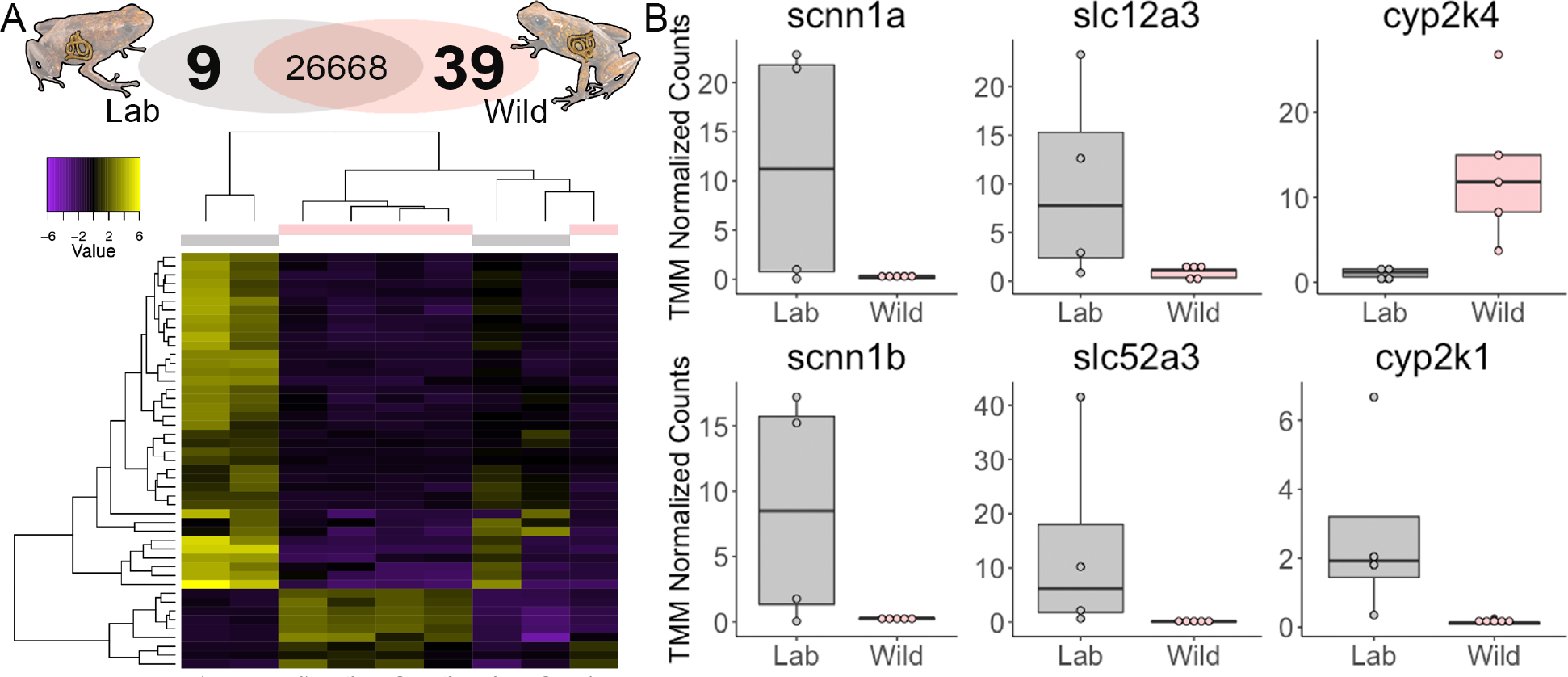
Intestinal gene expression differences between laboratory-reared and wild poison frogs. **(a)** Number of transcripts differentially expressed in captive reared (grey, N=4) and wild caught (pink, N=5) Little Devil frogs is shown on the top. The heatmap shows clustering of individual frogs (columns) and differentially expressed transcripts (rows, yellow is relative increased expression and purple is relative decreased expression). **(b)** Differentially expressed genes involved in sodium or small molecule transport or metabolism based on a modified t-test using DESeq2 and corrected for false discovery rate. This experiment was performed once. Gene name abbreviations: scnn1a, sodium channel epithelial 1 alpha subunit; scnn1b, sodium channel epithelial 1 beta subunit; slc12a3, solute carrier family 12 member 3; slc52a3, solute carrier family 52 member 3; cyp2k4, cytochrome P450 2K4; cyp2k1, cytochrome P450 2K1.

#### Skin

A total of 48 transcripts had significant differential expression, with chemically defended wild caught frogs having increased expression of 39 transcripts and decreased expression of 9 transcripts compared to the laboratory-raised frogs (Fig 3a). Only two GO enrichment terms survived false discovery rate correction, both with molecular function enrichment in RNA polymerase II function (p = 0.046). Of particular note, we found differential expression of the beta 2 subunit of the sodium potassium ATPase pump. Mutations in the alpha subunit of this ion pump are well known to be associated with chemical defenses in a number of animals that eat or sequester small molecules targeting this specific protein (Ujvari et al., 2015).

**Figure 3.**
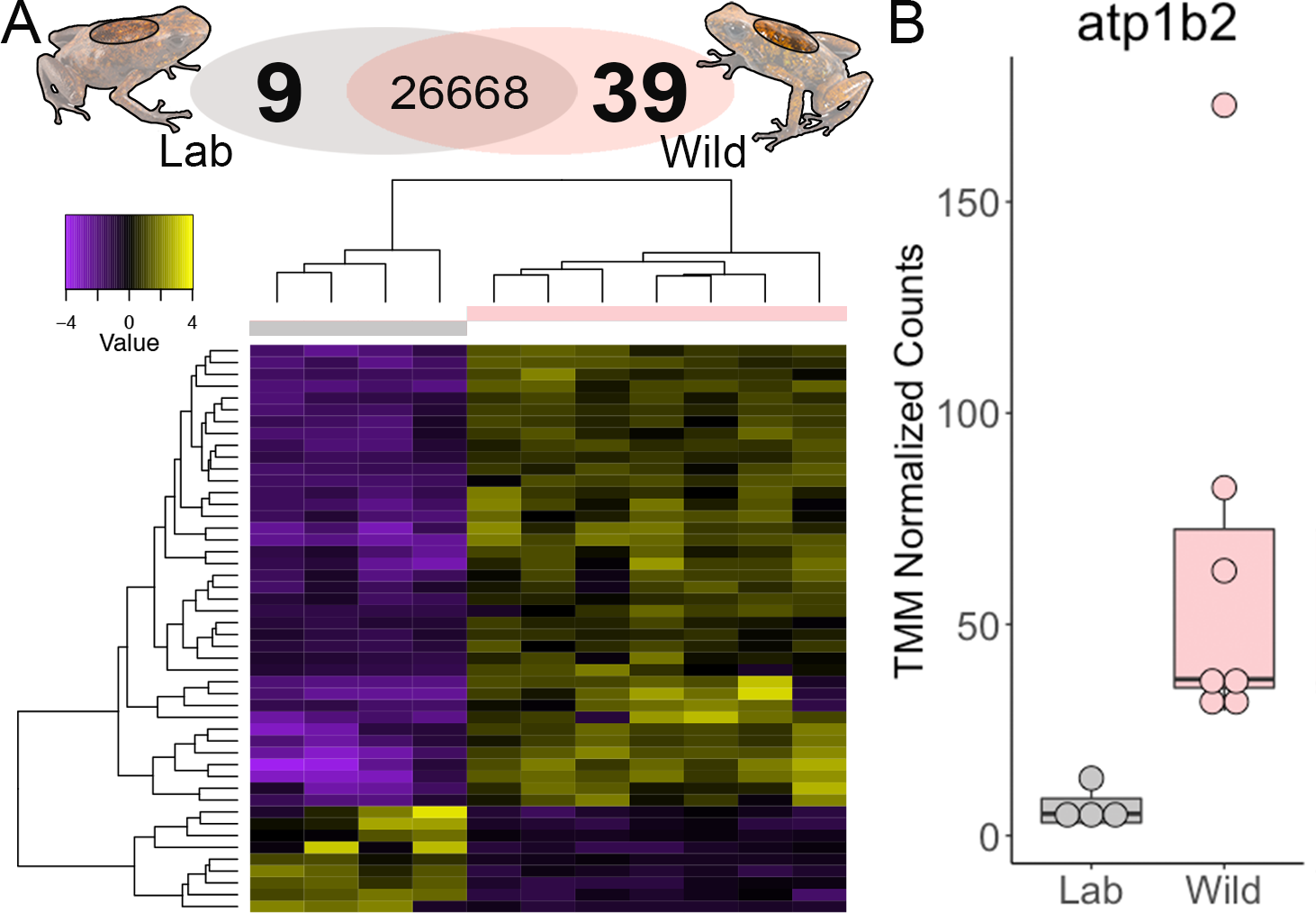
Skin gene expression differences between laboratory-reared and wild poison frogs. **(a)** Number of transcripts differentially expressed in captive reared (grey, N=4) and wild caught (pink, N=7) Little Devil frogs is shown on the top. The heatmap shows clustering of individual frogs (columns) and differentially expressed transcripts (rows, yellow is relative increased expression and purple is relative decreased expression). **(b)** Differentially expressed genes (based on a modified t-test using DESeq2 and corrected for false discovery rate) include the sodium-potassium-transporting ATPase subunit beta 2 (atp1b2), which is involved in sodium transport. This experiment was performed once.

#### Liver

In the liver, 75 transcripts had significant differential expression, with chemically defended wild caught frogs having increased expression of 54 transcripts and decreased expression of 21 transcripts compared to the laboratory-reared frogs (Fig 4a). Although there were more transcripts differentially expressed in the liver with better group clustering compared to the intestines (see heatmap in Fig 2a), no GO enrichment terms survived false discovery rate correction. Despite this, we noted several functional themes among the differentially expressed transcripts. There were several cytochrome p450s involved in small molecule metabolism, several small molecule transport proteins, including apolipoprotein A-IV and solute carrier proteins, and several molecular chaperones (heat shock proteins) (FIg 4b). Most of these transcripts involved in small molecule metabolism and transport had higher expression in the wild caught chemically defended frogs compared to the laboratory-reared frogs.

**Figure 4.**
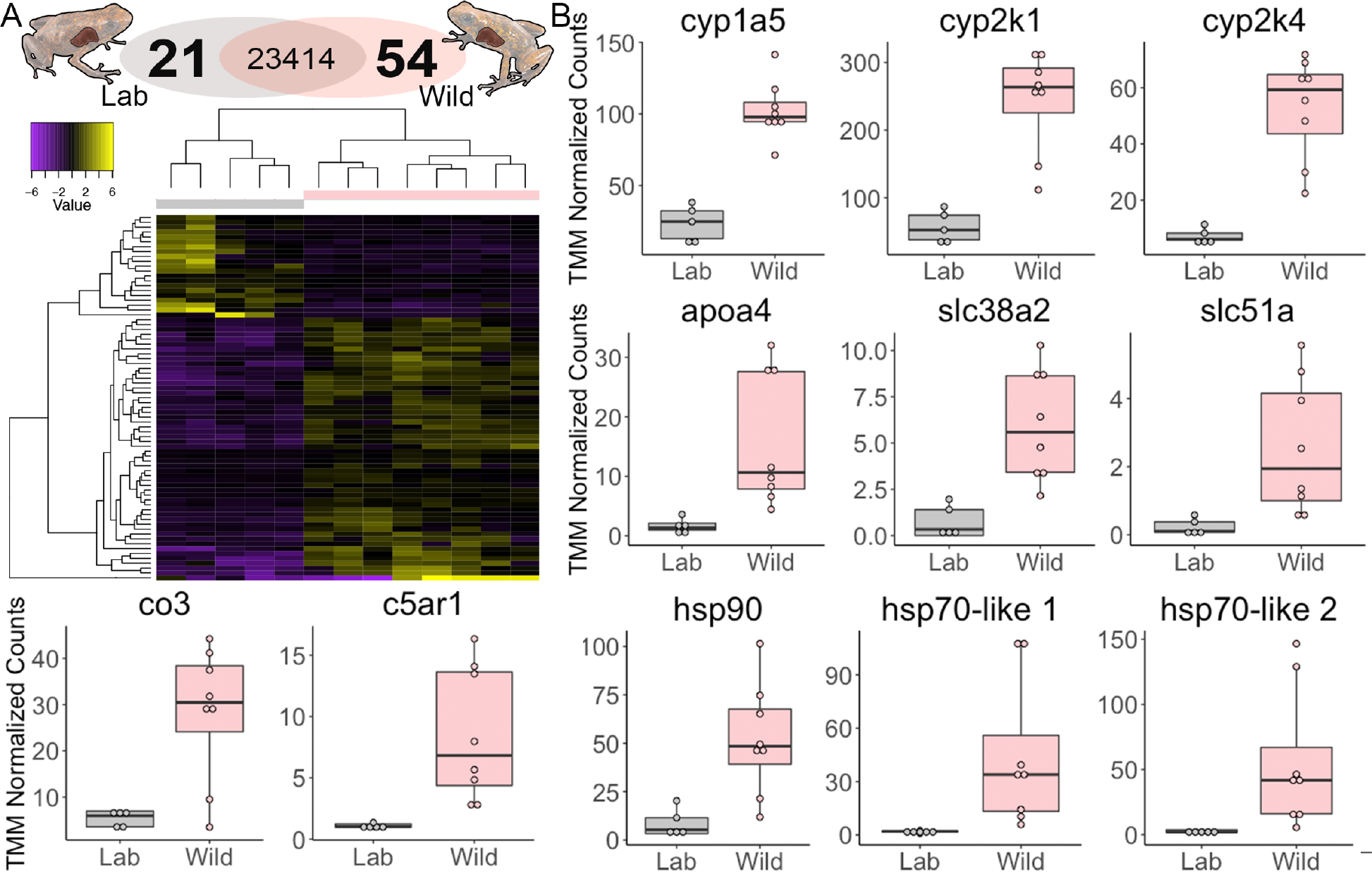
Liver gene expression differences between laboratory-reared and wild poison frogs. **(a)** Number of transcripts differentially expressed in laboratory-reared (grey, N=5) and wild caught (pink, N=8) Little Devil frogs is shown on the top. The heatmap on the bottom shows clustering of individual frogs (columns) and differentially expressed transcripts (rows, yellow is relative increased expression and purple is relative decreased expression). **(b)** Differentially expressed genes (based on a modified t-test using DESeq2 and corrected for false discovery rate) involved in small molecule transport or metabolism. This experiment was performed once. Gene name abbreviations: cyp1a5, cytochrome P450 1A5; cyp2k1, cytochrome P450 2K1; cyp2k4, cytochrome P450 2K4; apoa4, apolipoprotein A-IV; slc38a2, solute carrier family 38 member 2; slc51a, solute carrier family 51 member a; co3, complement C3; c5ar1, C5a anaphylatoxin chemotactic receptor 1; hsp90, Heat shock protein HSP 90-alpha; hsp70-like 1, heat shock 70 kDa protein 1-like; hsp70-like 2, heat shock 70 kDa protein 2-like.

### Proteomic analyses identify candidate toxin-binding proteins

We reasoned that plasma proteins may shuttle these lipophilic small molecules to their final destination. Therefore, we quantified plasma proteins in chemically defended wild caught frogs and laboratory frogs reared on an alkaloid-free diet. The most abundant plasma proteins identified were fetuins (fetuin-B and alpha-2-macroglobulins), apolipoproteins (A-I and A-IV), albumin, hemoglobin, transferrins (serotransferrin and saxiphilin), and immune-related proteins (complement factors and immunoglobulins). Ten proteins were identified as having significant differences in abundance between groups, although these did not survive false discovery adjustments, likely due to the small samples size (See Supplementary Data File for data and statistics). Among these proteins was saxiphilin (p=0.02), an amphibian transferrin-like protein that binds the neurotoxin saxitoxin (Morabito and Moczydlowski, 1995), which was more abundant in laboratory-reared frogs (Fig 5). Complement C3 was significantly higher in wild frogs compared to laboratory frogs raised on an alkaloid-free diet (p=0.04), supporting gene expression patterns in the liver (Fig 4b). Finally, plasma alpha-2-macroglobulin protein was also higher in chemically defended wild frogs compared to laboratory-reared frogs (p=0.05).

**Figure 5.**
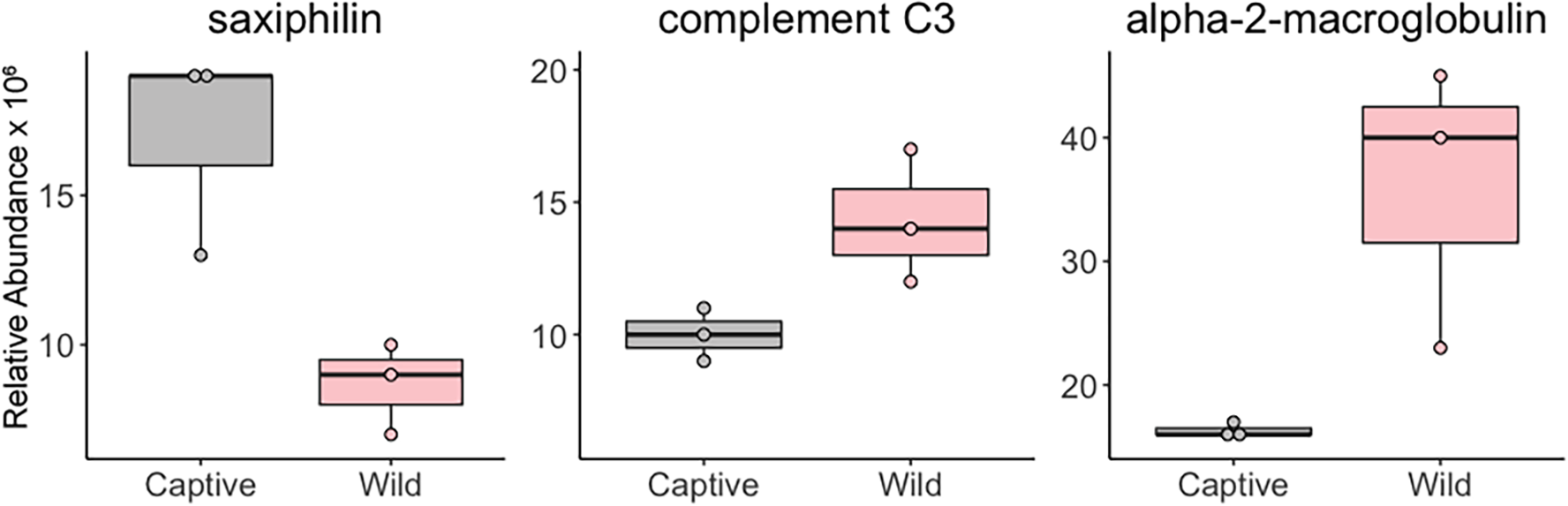
Plasma protein abundance differences between laboratory-reared and wild poison frogs. Laboratory-reared (grey, N=3) and wild caught (pink, N=3) Little Devil frogs have abundance differences of saxiphilin, complement C3, and alpha-2-macroglobulin proteins Differences were significant based on a modified t-test, but significance did not survive false discovery rate correction. This experiment was performed once.

The liver makes many of the proteins found in blood. To identify candidate toxin-binding proteins that may bind alkaloids for transport in the circulation, we used an untargeted thermal proteome profiling approach to identify proteins that interact with unlabeled small molecules (Fig 6a) (Franken et al., 2015). The small molecule used in this assay was decahydroquinoline, a commercially available alkaloid that we have detected in the wild caught frogs (present study) and in other *O. sylvatica* frog populations previously (McGugan et al., 2016). Using this assay, we identified several soluble proteins whose thermal profile shifted more than 4C that parallels other proteins of interest identified from above (Fig 6b). This includes saxiphilin, which was higher in the plasma of laboratory-reared frogs compared to wild frogs (Fig 5), and heat shock protein 90, which had higher transcript abundance in the livers of chemically defended wild frogs compared to laboratory frogs raised on an alkaloid-free diet (Fig 4b).

**Figure 6.**
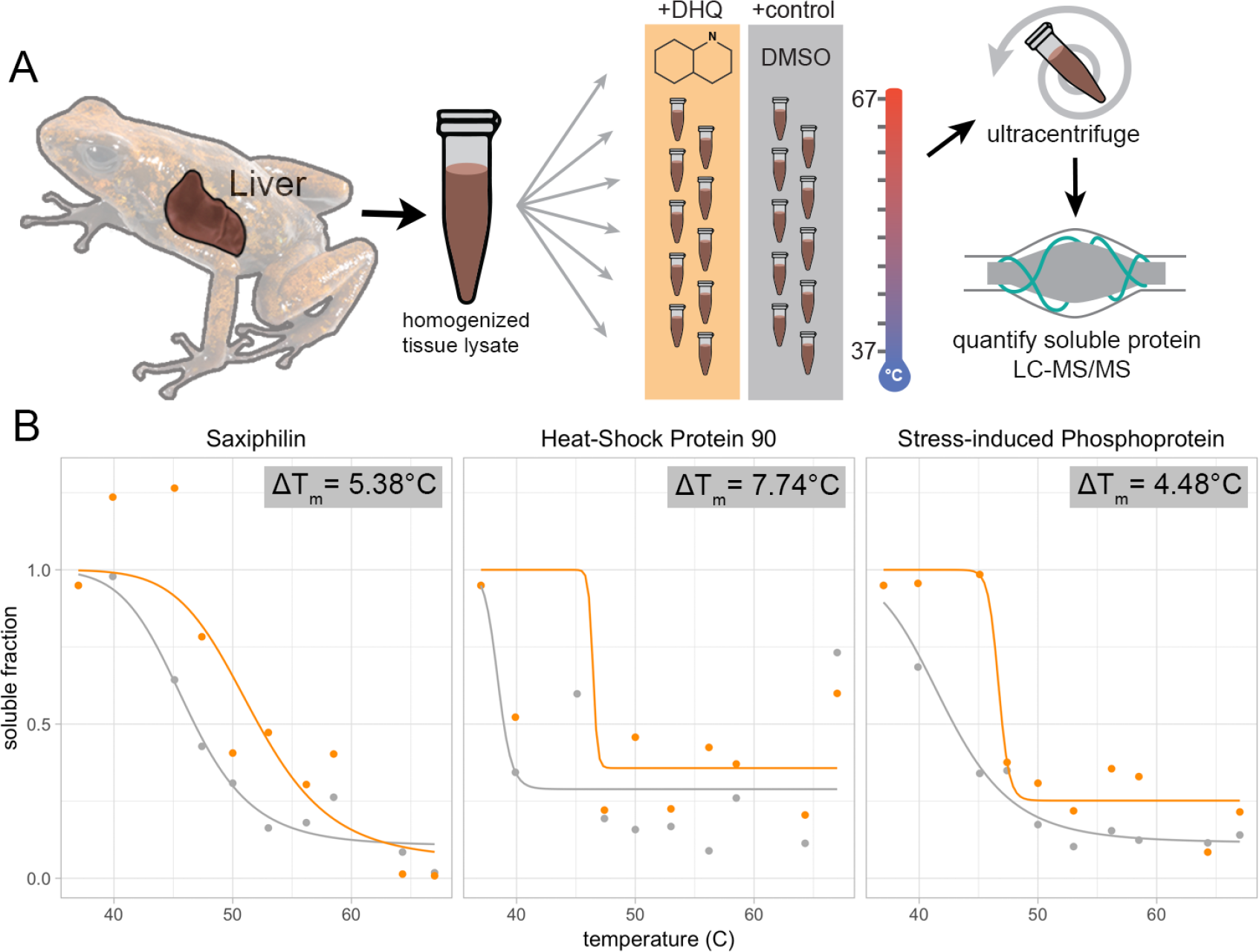
Thermal proteome profiling provides candidates for toxin-binding proteins in poison frogs. **(a)** Liver lysate from a single individual was treated with either decahydroquinoline (DHQ, orange) or DMSO (vehicle control, grey), aliquoted into 10 replicates and incubated at a range of temperatures prior to ultracentrifugation followed by quantitative proteomics using liquid chromatography tandem mass spectrometry (LC/MS-MS). **(b)** Examples of proteins with a greater than 4C shift in thermal stability when treated with DHQ (orange) compared to vehicle (grey). Melting temperature shifts are indicated in the top right of each graph; this experiment was performed once.

## Discussion

By comparing the physiology of wild caught chemically defended frogs to laboratory-raised frogs reared on an alkaloid-free diet, we were able to quantify differences in gene expression and protein abundance that may be involved in pathways linked to chemical defenses. In these comparisons, we found several themes in the differentially expressed transcripts in the intestines, liver and skin, as well as the plasma proteome, including involvement in small molecule metabolism. Moreover, several proteins were identified as possible candidate alkaloid-binding proteins across gene expression and ligand-binding experiments. Here we discuss these results in the context of possible physiological mechanisms of toxin sequestration and highlight the limitations of the present study with suggestions for future research priorities.

### Dietary alkaloid compounds

Many of the alkaloids identified in the Otokiki population were indolizidines and lehmizidines. These small molecules have been identified in other *O. sylvatica* populations and many other dendrobatid poison frogs (McGugan et al., 2016; Myers and Daly, 1976). All alkaloids described here have been noted in *O. pumilio* over 30 years of intense chemical ecology research (Saporito et al., 2007). Research in the 1970s identified histrionicotoxins as the major alkaloid group in *O. sylvatica* ((Myers and Daly, 1976), referred to as *D. histrionicus*). However, our present data from the Otokiki population, as well as previous work on three other *O. sylvatica* populations, did not include histrionicotoxins as the major alkaloid (McGugan et al., 2016), but rather lehmizidines and several classes of indolizidines. There could be several reasons for the absence of histrionicotoxins in *O. sylvatica* in this study. Myers and Daly (1976) sampled *O. sylvatica* in the southernmost part of their Ecuadorian range and did not examine northern populations (e.g. Otokiki); perhaps southern populations contain histrionicotoxins whereas northern Ecuadorian populations do not. Alternatively, since diet is a major component of poison frog alkaloid profiles (A. Saporito et al., 2009), arthropod diversity within the *O. sylvatica* range may have shifted in the last 40 years to chemically distinct prey items that make indolizidines the dominant alkaloid group. Population variation in toxin profiles is well established among various poison frogs species (Saporito et al., 2006)(Stuckert et al., 2014) and so broader sampling of many populations would reveal a more complete picture of dominant alkaloids in this species.

We detected zero to trace amounts of alkaloids in frogs moved from the wild as juveniles and raised in the laboratory for six months on a diet of crickets and fruit flies. This came as a surprise, as there are some reports of poison frogs retaining toxins for long periods of time in terrariums, including years for the Golden poison frog (*Phyllobates terribilis*) (Daly et al., 1980). It may be the age at which the laboratory frogs in our study were collected from the wild that influenced this result. Juvenile and subadult poison frogs tend to have a lower diversity and abundance of toxins in their skin [39] as well as smaller granular glands for storage [40]. It is possible that the laboratory frogs used in this study did not contain many alkaloids at the time they were collected and were then subsequently lost in the laboratory. It is also possible that there are species differences in alkaloid retention duration in captivity, although a robust toxin loss experiment had never been reported. It is also conceivable that different toxin classes (batrachotoxin versus indolizidines) are retained for varying amounts of time. For example, there were trace amounts of pumiliotoxin **307B** and some lehmizidines in the laboratory-reared frogs and perhaps these are retained for longer periods of time than indolizidines. The rate of alkaloid uptake and loss in adult poison frogs is understudied and such pharmacokinetics studies would lend insight into the dynamics of toxin sequestration and loss. Regardless, tissue samples from chemically defended and laboratory frogs collected from a single population enabled us to examine the physiology of alkaloid sequestration across various tissues.

### Sodium transporters

Sodium transport proteins were a common class of differentially expressed genes. In the intestines, we found significant downregulation of the epithelial sodium ion channel subunit transcripts (scnn1a and scnn1b; also called amiloride-sensitive sodium channel) in wild caught frogs compared to laboratory frogs reared on an alkaloid-free diet. This non-voltage sensitive sodium ion channel is inhibited by the diuretic alkaloid amiloride and is responsible for sodium influx across the apical membrane of intestinal epithelial cells. It is regulated by a number of intrinsic and extrinsic factors like diet and hormones (Bhalla and Hallows, 2008), which makes it difficult to determine how the various differences between the frogs groups studied here most influenced this result. Voltage gated-sodium channels are a well-known target of frog alkaloids (Santos et al., 2016), but to our knowledge the epithelial sodium channel has not been tested with poison frog alkaloids, making it unclear if this channel binds alkaloids. Alternatively, these channels could play a role in retaining alkaloids in the gut epithelium similar to resistance of tetrodotoxin in mantids (Mebs et al., 2016). Another transporter of note is the sodium-potassium-ATPase pump, whose beta 2 subunit expression was elevated in chemically defended wild caught frogs compared to laboratory frogs. The sodium-potassium-ATPase pump alpha subunit is a frequent target in the evolution of toxin resistance in a number of herbivorous invertebrates (Yang et al.)(Dobler et al., 2012) and vertebrates (Ujvari et al., 2015). Investigations of the function and evolution of the beta subunit in the context of toxin resistance is lacking. An analysis of sodium-potassium-ATPase pump sequence evolution across chemically defended and non-defended amphibians would likely yield interesting results. Binding studies with these channels and poison frog alkaloids as well as protein sequence evolution analyses are an important future goal in understanding potential interactions.

### Small molecule transport and metabolism

Vertebrates use two physiological mechanisms for transport of exogenous compounds depending on the degree of hydrophobicity: the portal venous system and the lymph system (Trevaskis et al., 2008). Cholesterol-derived bile acid molecules, which are made in the liver, are secreted to the gallbladder and then deposited into the intestines through the bile duct. With the help of bile acids and phospholipids, lipophilic molecules form micelles that are shuttled through the intestinal cells and excreted into the portal venous system. The transporter responsible for bile acid-associated export from intestinal cells into the portal blood is solute carrier protein 51a (slc51a, also known as organic solute transporter subunit alpha) (Dawson and Karpen, 2014), which has increased expression in the liver of chemically defended wild frogs compared to laboratory frogs. Bile acid-associated pathways are of particular interest, as bile acid derivatives have been observed in the skin of mantellid poison frogs (Clark et al., 2012), suggesting a bile acid-based transport system for some poison frog alkaloids. Alkaloids can be also be carried by a number of transport proteins in the blood circulation. One protein of particular interest that was increased in laboratory frogs compared to chemically defended wild frogs is saxiphilin. Saxiphilin is an amphibian specific transferrin that has high specificity for the alkaloid neurotoxin saxitoxin (Morabito and Moczydlowski, 1995). We also found evidence through thermal proteome profiling that saxiphilin may also bind alkaloids, like decahydroquinoline. Saxiphilin abundance in the blood may be influenced by alkaloid quantity, where the protein is upregulated when alkaloids are scarce in an effort to capture more molecules. Alternatively, saxiphilin could be shuttled to specific tissues once bound to an alkaloid, pulling saxiphilin out of plasma circulation in chemically defended frogs. Clearly more research on saxiphilin regulation and potential function as a general alkaloid carrier are warranted..

Not all molecules are transported in the blood however, as highly lipophilic exogenous compounds are transported by lipoproteins that move lipophilic cargo in chylomicron particles through the lymphatic system (Trevaskis et al., 2008). Apolipoproteins are required for chylomicron assembly and play some role in determining the types of compounds that are transported in their cargo. In our study, chemically defended frogs had increased expression of an apolipoprotein A-IV (apoa4) transcript in the liver compared to laboratory frogs reared on an alkaloid-free diet. We should also note that these changes in gene expression could also be due to differences in diet between our two groups, where wild frogs eat mostly ants and mites and laboratory frogs were fed *Drosophila* and crickets, which differ not only in alkaloid content but also in lipid and protein content (Bukkens, 1997). A more controlled diet experiment and additional biochemical testing would be required to completely understand the involvement of these candidate transport mechanisms in diet and chemical defenses of poison frogs.

Solute carrier proteins are of special interest as they aid in the transport of ions and small molecules across cell membranes, which is likely important for alkaloid transport in both the blood and the lymphatic system. In the intestines, two solute carrier family transcripts had decreased expression in wild caught frogs compared to laboratory-reared frogs. One of the transporters is slc12a3, a sodium and chloride cotransporter that is sensitive to a number of thiazide-like alkaloid diuretics that are used to treat hypertension (Gamba, 2009); the role of this transporter in the intestines is not well understood. The other differentially expressed transporter in the intestines is slc52a3, which transports riboflavin (vitamin B2, also an alkaloid) and is well-studied as loss of function due to mutations cause Brown-Vialetto-Van Laere syndrome (Yonezawa and Inui, 2013). In both cases, whether these proteins can transport other compounds is not known, although their expression can be regulated by diet, such as riboflavin intake (Said and Mohammadkhani, 1993). As these transporters interact with plant alkaloids naturally, it remains an open question as to whether poison frog alkaloids also influence their activity and gene expression. In the liver, the other small molecule transport protein differentially expressed between groups was slc38a2 (also called SNAT2), which had increased transcript abundance in wild frogs compared to laboratory frogs. This protein functions as a sodium-dependent amino acid transporter and its expression is heavily regulated by nutritional factors, where amino acid withdrawal induces slc38a2 expression *in vitro* (Hoffmann et al., 2018). As ants have lower amino acid content within a hard exoskeleton compared to softer fruit flies and crickets (McCusker et al., 2014), slc38a2 expression in these ant-specialist frogs is likely correlated with diet variation between frog groups rather than the presence of alkaloids.

Several cytochrome P450s transcripts were differentially expressed in the intestines and liver. This large protein family is well known for small molecule metabolism and has important clinical implications in drug metabolism and toxin breakdown (Danielson, 2002). We found three cytochrome P450s were differentially expressed between chemically defended wild caught frogs and laboratory-reared frogs. In both the intestines and liver, two transcripts in the cytochrome P450 2 family (cyp2k1-like and cyp2k4-like) were differentially expressed between groups. These enzymes were originally identified in rainbow trout (Buhler et al., 1994) and their specific chemical function has not been investigated in detail. In the liver, a cytochrome P540 1A enzyme (cyp1a5-like) was also differentially expressed between groups. This enzyme is well described in birds and is located in liver microsomes where it oxidizes a number of structurally unrelated compounds, including xenobiotics and steroids (Kubota et al., 2008; Shang et al., 2013). It is important to note that many biological factors can influence the expression of cytochrome P450s including diet, sex, and temperature (Haasch, 2002). We cannot conclude based on this study design if the presence of alkaloids induced these changes in gene expression and further experiments would be necessary to determine the dynamic regulation of these important enzymes. However, some poison frog species are thought to metabolize alkaloids, where John Daly and colleagues hypothesized a yet unidentified “pumiliotoxin 7-hydroxylase” was responsible for the metabolism of PTX **251D** into the more potent aPTX **267A** (Daly et al., 2003). Moving forward, a better understanding of alkaloid metabolism is needed in order to link poison frog alkaloid profiles to metabolism by cytochrome P450s.

### Other proteins classes of interest

Two other classes of proteins were different in many of the comparisons between wild caught chemically defended frogs and laboratory-reared frogs described here: proteins involved in immune system function and heat shock proteins. Two proteins in the complement immune system showed differences between groups. Complement C3 had increased transcript abundance in the liver and higher protein abundance in the blood in wild chemically defended frogs compared to laboratory-reared frogs. The membrane bound C5a receptor also showed higher transcript abundance in livers of wild frogs compared to laboratory frogs. Although complement factors can bind alkaloids (Garcia et al., 2017), abundance can also be influenced by pathogens and other environmental factors (Gasque, 2004). Thus, further experiments with more controlled groups are necessary to determine if these immunity factors are important for chemical defenses. The other class of molecules with differences in transcript abundance between groups was liver heat shock proteins. Although expression of these proteins can be modulated by stress and environmental variables like temperature (Feder and Hofmann, 1999), we also found evidence that heat shock protein 90 may bind the poison frog alkaloid decahydroquinoline. Other alkaloids have also been found to both decrease and increase the function of the heat shock protein chaperones (Wisén and Gestwicki, 2008) and, therefore, it is possible that poison frog alkaloids also modify heat shock protein activity. Alternatively, the alkaloids themselves could destabilize liver lysate proteins which then leads to an increased thermal shift of heat shock proteins in the thermal proteome profiling assay, where heat shock proteins are binding to proteins as they begin to unfold in the presences of decahydroquinoline. More thorough *in vitro* studies are needed to test these hypotheses.

### Conclusions and future directions

Our study has provided many candidate proteins and biological processes that may be involved in alkaloid sequestration in poison frogs. This includes candidate mechanisms for alkaloid transport like plasma carrier proteins and chylomicrons as well as genes involved in small molecule metabolism. However, a limitation of our study is that we compared gene expression between chemically defended wild caught frogs and frogs reared on an alkaloid-free diet that were in different environments. Future studies should be conducted in identical environmental conditions with controlled toxin feeding regimes. Finally, although our thermal proteome profiling experiment highlighted candidate proteins, this method exclusively tests soluble proteins and further studies should be done to test membrane bound proteins as well. Overall, we have provided the first organismal perspective into how poison frogs bioaccumulate small molecule chemical defenses from their diet. This work provides a foundation for future mechanistic and comparative work that will be able to uncover how poison frogs have altered their physiology to acquire chemical defenses.

### Data Availability

All mass spectrometry data (GC/MS data from the alkaloid analysis as well as LC-MS/MS files from plasma proteomics and thermal proteome profiling experiments), and the *O. sylvatica* transcriptome and proteome fasta files are available at DataDryad (submission pending). All Illumina fastq files are available on the Sequence Read Archive (submission pending). An excel file with all data (gene expression and peptide counts as well as alkaloid quantification) and statistical output is available in supplementary materials.

## Acknowledgements

We would like to thank Lola Guarderas (Wikiri) for support in Ecuador, Adam Freedman and Aaron Kitzmiller for advice on analyses, and Eugenia Sanchez for comments on early versions of this manuscript. All computational work was performed on the Odyssey cluster supported by the FAS Science Division Research Computing Group at Harvard University.

## Author Contributions

LAO and LAC designed the research; LAO, SNC, ABR, EET and LAC collected frog samples in the Ecuador; EET and LAC maintained the laboratory frogs; LAO performed the laboratory research and RNAseq analysis; GDB and SAT performed the small molecule mass spectrometry; GDB and CV performed the alkaloid GC/MS analysis; BB conducted the plasma proteomics experiments; AAB performed the thermal proteome profiling analysis; SNC and LAO wrote the paper with contributions from all authors.

